# Altered intrinsic ignition dynamics linked to Amyloid-*β* and tau pathology in Alzheimer’s disease

**DOI:** 10.1101/2024.03.29.587333

**Authors:** Gustavo Patow, Anira Escrichs, Noelia Martínez-Molina, Petra Ritter, Gustavo Deco, Alzheimer’s Disease Neuroimaging Initiative

## Abstract

Alzheimer’s disease (AD) progressively alters brain structure and function, yet the associated changes in large-scale brain network dynamics remain poorly understood. We applied the intrinsic ignition framework to resting-state functional MRI (rs-fMRI) data from AD patients, individuals with mild cognitive impairment (MCI), and cognitively healthy controls (HC) to elucidate how AD shapes intrinsic brain activity. We assessed node-metastability at the whole-brain level and in 7 canonical resting-state networks (RSNs). Our results revealed a progressive decline in dynamical complexity across the disease continuum. HC exhibited the highest node-metastability, whereas it was substantially reduced in MCI and AD patients. The cortical hierarchy of information processing was also disrupted, indicating that rich-club hubs may be selectively affected in AD progression. Furthermore, we used linear mixed-effects models to evaluate the influence of Amyloid-*β* (A*β*) and tau pathology on brain dynamics at both regional and whole-brain levels. We found significant associations between both protein burdens and alterations in node-metastability. Lastly, a machine learning classifier trained on brain dynamics, A*β*, and tau burden features achieved high accuracy in discriminating between disease stages. Together, our findings highlight the progressive disruption of intrinsic ignition across whole-brain and RSNs in AD and support the use of node-metastability in conjunction with proteinopathy as a novel framework for tracking disease progression.

## 1 Introduction

Among neurodegenerative disorders, Alzheimer’s Disease (AD) represents the most prevalent cause of dementia, initially affecting the medial temporal lobe (MTL) and the limbic system before gradually spreading to associative and primary neocortical regions^1–3^. Neurodegeneration, defined as the progressive deterioration of neuronal structure and function, can be found in the brain at many different organizational levels, from molecular to whole-brain systems^4^. Early symptomatic stages of AD are often captured under the clinical category of mild cognitive impairment (MCI), characterized by objective cognitive decline with minimal impact on daily living. As the disease progresses, deficits extend to multiple cognitive domains, including language, visuospatial orientation, and executive function, culminating in dementia due to AD^5^,^6^. The associated clinical burden significantly diminishes quality of life for patients and caregivers and imposes substantial socioeconomic costs worldwide^7^. Despite advances in diagnostic frameworks, including biomarker-based criteria, current diagnostic tools still suffer from limited sensitivity and specificity, with reported misdiagnosis rates of up to 20%^8^. Consequently, there is an urgent need for robust, noninvasive biomarkers that can capture early-stage neurophysiological alterations and monitor disease progression with high accuracy.

The clinical diagnosis of AD is primarily based on physiological biomarkers, including the presence of Amyloid-*β* (A*β*) plaques, alterations in cerebrospinal fluid, p-tau, and structural atrophy in MTL, all of which correspond to the ATN framework (Amyloid, Tau, Neurodegeneration)^9^,^10^. Recent neuroimaging studies using AD’s resting-state fMRI (rs-fMRI) data, however, have revealed stage-dependent brain activity fluctuations in canonical resting-state networks (RSNs) such as the default mode network, salience network, dorsal attention network, and limbic network (LN)^11–19^. From a dynamic systems perspective, metastable neuronal dynamics are reduced in the AD spectrum in association with disrupted network topology suggesting that measures of brain dynamics reflect important information of disease progression^20^. In light of this, one intriguing avenue to investigate is whether the intrinsic capability of the brain to integrate information over the whole-brain network, as measured with node-metastability, could serve as a potential biomarker.

Neuroscience frameworks grounded in complex systems theory, such as the Intrinsic Ignition Framework (IIF)^21^, allow quantifying dynamical properties essential for efficient information processing and can be applied at regional and whole-brain levels. Intrinsic ignition refers to the capacity of brain regions to initiate and sustain neural events that can propagate throughout the whole-brain network, as assessed by measures of node-metastability^22^. Node-metastability, in this context, indicates the local degree of functional variability of each brain area over time. Remarkably, this framework has demonstrated robust sensitivity in capturing differences in whole-brain dynamics across different brain states, encompassing both health and disease, such as deep sleep, meditation, aging, depression, and atypical development^23–27^.

In this study, we set out to investigate the effects of AD on brain dynamical complexity, understood as the broadness of communication, using the IIF^21^ applied to rs-fMRI data. Specifically, we examined global and RSN node-metastability in 36 subjects from the Alzheimer’s Disease Neuroimaging Initiative (ADNI3) across three disease stages: healthy controls, individuals with MCI, and patients with AD. Furthermore, we employed multilevel statistical modeling to assess the impact of proteinopathy —namely, Amyloid-*β* and tau protein burden— on node-metastability. We hypothesized that proteinopathy substantially decreases the node-metastability at both whole-brain and RSN levels. Finally, we performed machine learning classification to evaluate whether features derived from the IIF and protein burden could accurately discriminate between clinical stages. We hypothesized that node-metastability, in combination with protein burden, could offer a sensitive and specific framework for clinical stage classification.

## 2 Methods

### 2.1 Participants

Empirical data were obtained from the Alzheimer’s Disease Neuroimaging Initiative (ADNI3) database (adni.loni.usc.edu). The ADNI was launched in 2003 as a public-private partnership, led by Principal Investigator Michael W. Weiner. For up-to-date information, see www.adni-info.org.

In this study, we included a cohort comprising 17 healthy controls (HC), 9 individuals with mild cognitive impairment (MCI), and 10 patients with Alzheimer’s Disease (AD). These are the same participants as reported in^28^ and substantially overlap with those analyzed in Stefanovski et al.^29^ and Triebkorn et al.^30^. As an inclusion criterion for AD patients, the diagnosis criteria of NINCDS-ADRDA from 1984 were used, which contains only clinical features^6^. Inclusion criteria for both HC and MCI were a Mini-Mental State Examination (MMSE) score between 24 and 30 and age between 55 and 90 years. Additionally, for MCI, the participant must have a subjective memory complaint and abnormal results in another neuropsychological memory test. The MMSE score had to be below 24 to fulfill the criteria for AD, and the NINCDS-ADRDA criteria for probable AD had to be fulfilled^6^. Imaging and biomarkers were not used for the diagnosis. For the full inclusion criteria of ADNI-3, see the study protocol [here](http://adni.loni.usc.edu/wp-content/themes/freshnews-dev-v2/documents/clinical/ADNI3_Protocol.pdf).

To ensure adequate statistical power, we performed an a priori sample size calculation using G *\** Power^31^, specifying a two-group Wilcoxon-Mann-Whitney test, with a significance level of *α* = 0.05 and power 1*− β* = 0.8. Assuming a standard deviation of *σ* = 0.05 — empirically justified by our observed data—, this design permits detection of an effect size of at least Cohen’s *d* = 1.1.

### 2.2 Data Acquisition

All images used in this study were downloaded from ADNI3. To allow comparisons, only data from Siemens scanners with a magnetic field strength of 3T were used (models: TrioTim, Prisma, Skyra, Verio). For full details regarding acquisition parameters, please see Supplementary Material in ^29^.

We included the following imaging modalities: T1-weighted (MPRAGE) images, TE = 2.95–2.98ms, TR = 2.3 s, matrix and voxel size differ slightly; FLAIR images, TE differs slightly, TR= 4.8 s, matrix size = 160 *×* 256 *×* 256, and voxel size differs slightly; rs-fMRI T2*-weighted echo planar images, TE=30ms, TR= 3s, matrix size = 64 *×* 64 *×* 48, and voxel size = 3.4 mm^3^, 197 volumes; Siemens Fieldmaps and PET images (AV-45 for A*β* and AV-1451 for tau).

Imaging data preprocessing can be subdivided into structural images, rs-fMRI, and PET. Here we provide a brief description of the preprocessing pipelines, for more details please see ^28^.

### 2.3 MRI preprocessing and brain parcellation

For each included participant, we created a brain parcellation using T1w, fieldmaps, and FLAIR images. We followed the minimal preprocessing pipeline^32^ of the Human Connectome Project (HCP) using [Freesurfer](https://surfer.nmr.mgh.harvard.edu/fswiki/FreeSurferMethodsCitation) ^33^, FSL^34–36^ and [Connectome Workbench](https://www.humanconnectome.org/software/connectome-workbench). We then registered the subject cortical surfaces to the parcellation of Glasser et al.^37^ using the multimodal surface matching (MSM) tool^38^. We mapped the parcellation on the surface back into the gray matter volume with the connectome workbench. Our parcellation included 379 regions: 180 left and 180 right cortical regions, 9 left and 9 right subcortical regions, and 1 brainstem region.

We examined differences in brain dynamics between HC, MCI, and AD subjects in the seven Resting State Networks (RSNs), included in the Schaefer parcellation^39^. To transfer the RSNs from the original parcellation to the parcellation of Glasser et al.^37^, we used the 1000-node version of the Schaefer parcellation. We transferred the respective RSN label of the closest node in the Schaefer parcellation for each node in the target parcellation, using the Euclidean distance as a measure.

### 2.4 rs-fMRI preprocessing

The preprocessing of rs-fMRI was computed using FSL FEAT and independent component analysis–based denoising (FSLFIX), an independent component analysis–based denoising method that uses an automated classifier to identify noise-related components for removal from the data. We used a standard pipeline^29^, which included removal of the first four volumes, rigid-body head motion correction, 3-mm spatial smoothing to improve signal-to-noise ratio, and a high-pass temporal filter of 75s to remove slow drifts. The data were then denoised using FSLFIX. The algorithm was trained on a manually labeled held-out set of 25 individuals scanned with identical imaging parameters. Using ordinary least squares regression, time courses for noise-labeled components and 24 head motion parameters were removed from the voxel-wise fMRI time series.

The resulting denoised functional data were spatially normalized to Montreal Neurological Institute (MNI) space using Advanced Normalization Tools (version 2.2.0), via a three-step method: (I) registration of the mean realigned functional scan to the skull-stripped high-resolution anatomical scan via rigid-body registration; (II) spatial normalization of the anatomical scan to the ICBM152 MNI template via a non-linear registration; and (III) normalization of the functional scan to the MNI template using a single transformation matrix that concatenates the transforms generated in steps I and II. Mean time series for each parcellated region were then extracted and filtered in the range 0.01 to 0.09 Hz.

### 2.5 PET preprocessing

For A*β*, we used the preprocessed version of AV-45 PET by ADNI. These images included the following preprocessing steps: Images acquired 30–50 min post tracer injections: four 5-min frames (i.e., 30–35 min, 35–40 min…). These frames are co-registered to the first and then averaged. The averaged image was linearly aligned such that the anterior-posterior axis of the subject is parallel to the AC-PC line. This standard image has a 1.5 mm cubic voxel resolution and a matrix size of 160 *×* 160 *×* 96. Voxel intensities were normalized so that the average voxel intensity was 1. Finally, the images were smoothed using a scanner-specific filter function. The filter functions were determined in the certification process of ADNI from a PET phantom. We used the resulting image and applied the following steps. First, the PET image must be rigidly aligned with the participant’s T1 image (after being processed in the HCP structural pipeline). The linear registration was done with FLIRT (FSL). The PET image was then masked with the subject-specific brainmask derived from the structural preprocessing pipeline (HCP). To obtain the local burden of A*β*, we calculated the standardized uptake value ratio (SUVR) with the cerebellum as a reference region because it is known that the cerebellum does not show relevant AV-45 PET signals ^40^,^41^. We therefore receive in each voxel a relative A*β* burden. The cerebellar white matter mask was taken from the Freesurfer segmentation (HCP structural preprocessing). The image was then partial volume corrected using the Müller-Gärtner method from the PETPVC toolbox ^42^. For this step, the gray and white matter segmentation from Freesurfer (HCP structural preprocessing) was used. Subcortical region PET loads were defined as the average SUVR in subcortical GM. Cortical GM PET intensities were mapped onto the individual cortical surfaces using the connectome workbench tool with the pial and white matter surfaces as ribbon constraints. Using the multimodal parcellation from ^37^, we derived average regional PET loads.

For tau, we used the ADNI-preprocessed AV-1451 PET (flortaucipir) images, which followed an identical acquisition and preprocessing pipeline. This resulted in a single relative tau value per voxel, which was similarly averaged within each region of the Glasser parcellation.

### 2.6 Intrinsic Ignition Framework (IIF)

We used the IIF^21^,^43^to examine the dynamical complexity across three groups (HC, MCI, and AD). The framework assesses the degree of whole-brain integration based on spontaneously occurring events over time. The methodology for calculating intrinsic integration values across brain areas is illustrated in **Figure 1**. The algorithm involves identifying driving events for each brain area, which are converted into a binary signal using a threshold^44^. To represent events as a binary signal, the time series is transformed into z-scores, denoted as *z*_*i*_(t), and a threshold value, *θ*, is applied. Specifically, an event is marked as 1 in the binary sequence *σ* (t) if *z*_*i*_(t) surpasses the threshold from below and marked as 0 otherwise. Upon triggering an event, the neural activity is measured in all brain areas within a time window of 4TRs. This window width was selected by the time it takes the integration to return to basal values. Then, a binary matrix is constructed to depict the connectivity between brain areas exhibiting simultaneous activity. The measure of global integration^45^ is then computed to assess the broadness of communication across the network for each driving event (i.e., the largest subcomponent). This process is iterated for each spontaneous neural event to obtain the node-metastability, quantified as the standard deviation of the integration for each brain area in the brain network. We computed the framework across the whole-brain network and within 7 large-scale RSNs (control, DMN, dorsolateral attention, limbic, somatomotor, salience, and visual networks).

**Figure 1.**
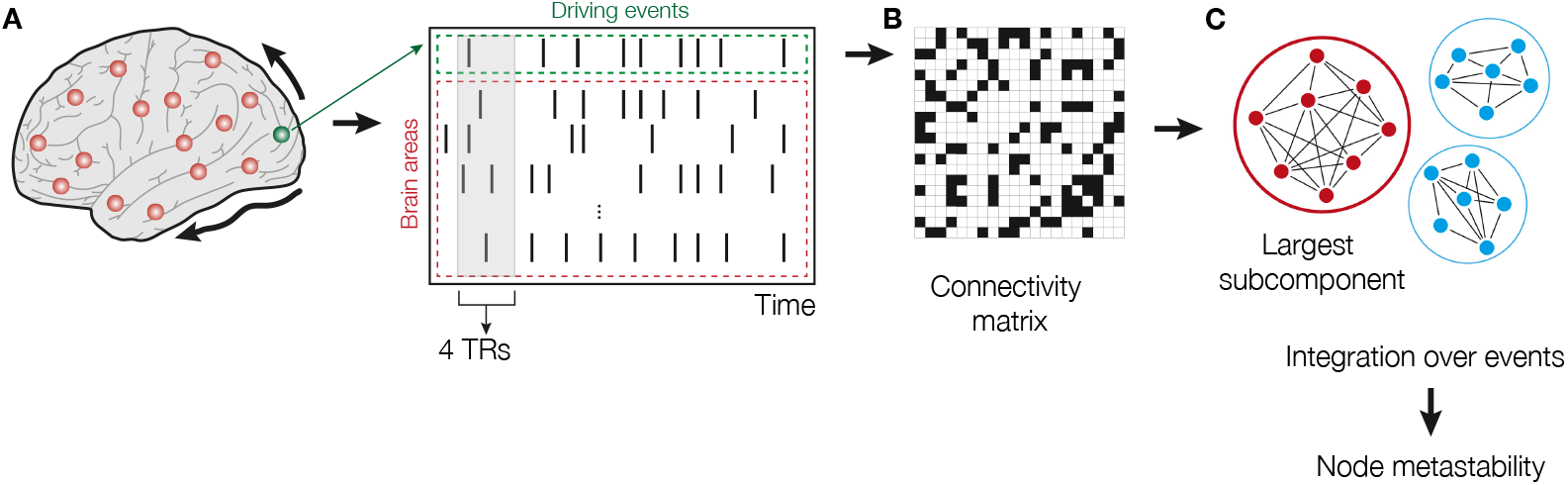
Intrinsic Ignition Framework. **(A)** Events were captured applying a threshold method^44^. For each driving event in a given brain area (green dashed area), the functional connectivity with the rest of the network was measured in the time window of 4TRs (red dashed area). The selection of this window width was determined by the time that takes the integration to return to basal values. **(B)** A binarized matrix was obtained, representing the connectivity between brain areas where activity was simultaneous. **(C)** Applying the global integration measure^45^, we obtained the largest subcomponent. Repeating the process for each driving event, we calculated the node-metastability computed as the standard deviation of the integration of each brain area over time. Figure modified from^24,25,43^.

We have also defined the Hierarchy Disruption Factor (HDF), which measures the *l*^2^ norm between two hierarchies, as:

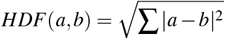

where *a* and *b* are two cohort-based (i.e., HC, MCI, or AD) hierarchies. A hierarchy for a cohort is defined by computing, for each node in the parcellation, the averaged metastability over all subjects in that cohort, and then sorting from largest to smallest values^43^. See Figure 2B.

**Figure 2.**
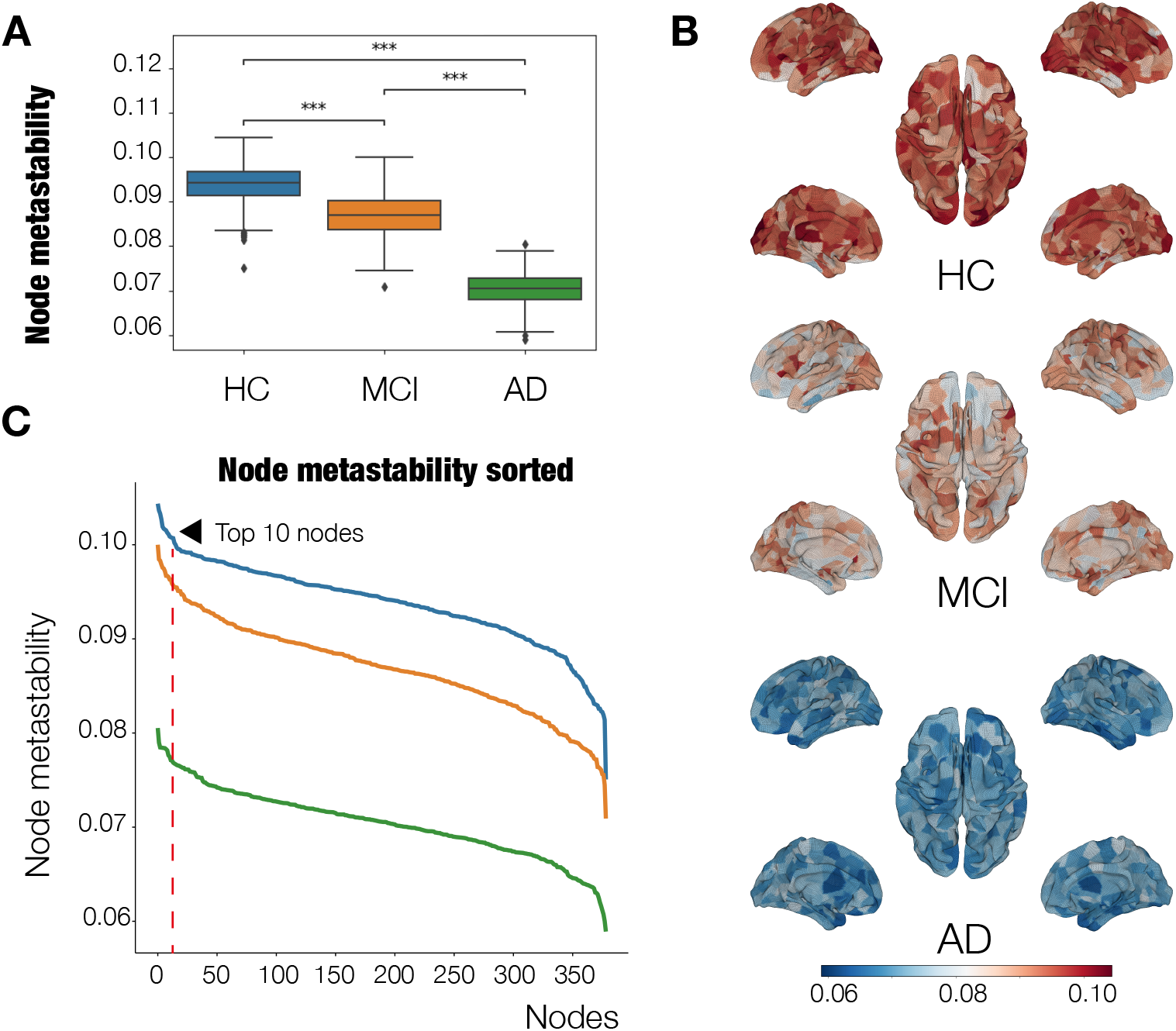
Dynamical complexity of Alzheimer’s Disease stages. **(A)** Node-metastability. HCs showed higher node-metastability values at the whole-brain level compared to MCI and AD patients. P-values are based on a permutation test, where *** represents *p <*= 1.00*e−* 04. **(B)** Hierarchy. The red line denotes the 10 regions showing the highest node-metastability values in each AD stage. For the HCs, brain areas showing the highest values were primarily located in the VIS, SM, and DAT networks. For the MCI stage, the brain areas belonged to the same networks, in addition to the VIS network, with a lower metastability. At the AD stage, they were located again in the VIS, SM, and DAT networks, although with a severe decrease in metastability. **(C)** Brain renders represent the node-metastability values of the 379 areas for each AD stage. The dynamical complexity of the HCs across the whole-brain networks is more complex than the dynamical complexity of the other two stages.

### 2.7 Statistical Analyses

Permutation tests were used for all group comparisons by comparing the empirical vs. the randomized data. To do so, we used a Monte Carlo permutation method. We randomly shuffled the labels between conditions to obtain two new simulated conditions (10,000 permutations). Then, we evaluated how much the difference between the simulated conditions was higher than between the actual conditions. We computed the p-value of the null hypothesis that the two random distributions show higher differences than the actual conditions. To correct for multiple comparisons, we used the Benjamini-Hochberg correction to control for false discovery rate (FDR).

### 2.8 Multilevel Modeling of AD on Resting State Networks

We implemented a Linear Mixed Effects model (LME) using the *lmer* function in R Statistical Software (v4.3.2)^46^ with the lme4 package^47^.

We first assessed whether the outcome variable (i.e., each node’s metastability at the whole-brain level) shows a significant change by specifying the node’s A*β* and tau levels, and their interaction, as fixed effects, and the participant ID (subject) as a random effect. Please note that we only have 3 levels for the cohort, thus, it is not recommended to include it as a random effect^48^. We defined the syntax for the model as *Meta ∼ ABeta*Tau* + (1|*ID*).

Next, to refine our analysis and mirror the studies performed in the rest of the paper, we repeated the previous analysis considering the node-metastability in the 7 RSNs as a grouping variable, both as a random and as a fixed effect. The final syntax for this model is *Meta ∼ ABeta*Tau*RSN −subcor* + (1 + *RSN −subcor*|*ID/RSN −subcor*).

In these analyses, we opted not to include the MMSE scores in the model, as this resulted in a more complex model without improving prediction power, as determined with pairwise ANOVA tests.

### 2.9 Machine learning analysis

We used machine learning (ML) analysis to test whether disease stage (HC, MCI, AD) could be classified using the brain dynamics (ignition, node-metastability) and proteinopathy (Amyloid*β* and tau SUVR) features. The k-nearest neighbors (KNN) classification algorithm was used to dissociate the diagnostic cohorts in this study (HC vs MCI, HC vs AD). The initial set of features included 1440 features per participant, which were the brain dynamics and proteinopathy features extracted from each of the 360 Glasser parcels. Data dimensionality was reduced by applying the feature selection algorithm minimum-Redundancy-Maximum-Relevance (mRMR)^49^ with a fixed number of 10 features to be selected to ensure comparability across classification analyses.

To enhance generalizability and mitigate overfitting, we adopted a nested cross-validation scheme with leave-one-out cross-validation (LOOCV) in the outer loop. This approach benefits small datasets, as it maximizes the training data used in each iteration. The inner loop of the nested structure was responsible for hyperparameter tuning — in this case, the number of neighbors (k) evaluated in the range 2 *< k <* 5. Feature selection (mRMR) was performed strictly within the training folds to prevent data leakage. After hyperparameter tuning, the best model was selected. As mentioned, in the outer loop, model evaluation was assessed on held-out participants, ensuring an unbiased estimation of generalization error.

In the inner loop, the dataset was split such that 75% of the data was used for training and 25% for validation. Stratified sampling was used to preserve the proportion of diagnostic classes across folds. Importantly, all pre-processing steps, including feature selection and hyperparameter tuning, were confined to the training data in each fold. The ML pipeline was implemented using Python 3.9, with core components from the Scikit-learn library ^50^, alongside auxiliary packages such as numpy, pandas, and shap for interpretability.

This methodological approach ensures that the reported classification accuracies reflect genuine predictive performance and are not inflated by circular analysis or overfitting. See Table 3.

## 3 Results

### 3.1 Clinical demographics

We included a cross-sectional cohort of 36 ADNI3 (adni.loni.usc.edu) participants: 17 healthy controls (HC), 9 individuals with mild cognitive impairment (MCI), and 10 patients with Alzheimer’s disease (AD). Group demographics, neuropsychological, and protein burden information are summarized in Table 1.

**Table 1.** Demographic, neuropsychological and protein burden information. Abbreviation: MMSE, mini-mental state examination. Diagnosis n (female) Mean *σ* Min. Max. Mean *σ*_*MMSE*_ Min. Max. Mean Mean

| Diagnosis | n (female) | Mean age | $\sigma$ | Min. age | Max. age | Mean MMSE | $\sigma_{MMSE}$ | Min. MMSE | Max. MMSE | Mean $A\beta$ | Mean tau |
| --- | --- | --- | --- | --- | --- | --- | --- | --- | --- | --- | --- |
| HC | 17 (10) | 70.8 | 4.3 | 63.1 | 78.0 | 29.3 | 0.7 | 28 | 30 | 1.31 | 1.53 |
| MCI | 9 (3) | 68.8 | 5.8 | 57.8 | 76.6 | 27.4 | 1.5 | 25 | 30 | 1.52 | 1.80 |
| AD | 10 (5) | 72.0 | 9.6 | 55.9 | 86.1 | 21.3 | 6.8 | 9 | 30 | 2.01 | 2.46 |

### 3.2 Node-metastability is attenuated at whole-brain level at MCI and AD stages

We computed the node-metastability measure to study the dynamical complexity underlying the whole-brain functional network in the three diagnostic groups, i.e., HC (0.094 *±* 0.0044, mean ± s.d.), MCI (0.087 *±* 0.0049), and AD (0.070 *±*0.0037). We found that the node-metastability significantly decreased in the AD group compared to the MCI (*P*_FDR-corr_ *<* 0.001, effect size *d* = 3.80) and HC groups(*P*_FDR-corr_ *<* 0.001, effect size *d* = 5.79). Furthermore, we found that the node-metastability was higher in HC than in MCI (*P*_FDR-corr_ *<* 0.001), effect size *d* = 1.45). Noteworthily, in all the cases, the difference between the average values is larger than the calculated minimum effect size we established before, of *d* = 1.1, given our sample size.

**Figure 2A** shows the results of this analysis, where we can see a clear, statistically significant difference between the three groups. The observed overall dynamical complexity systematically decreases as the disease progresses, which is to be expected as the different regions have their dynamics altered. **Figure 2B** shows the rendered brains representing the node-metastability for each group across the whole-brain functional network.

To supplement this analysis, we also computed the intrinsic ignition as the mean of the integration value, i.. the mean of the largest subcomponent calculated from the binarized matrix considered an adjacency matrix across nodes and time windows. Both MCI and AD exhibited a significant reduction in ignition compared to HC (*P*_FDR-corr_ *<* 0.001, see FigS1).

### 3.3 Hierarchy of information processing at whole-brain level appears disrupted at MCI and AD stages

**Figure 2C** shows the hierarchy for each disease stage across the whole-brain functional network (i.e., the sorted brain areas from highest to lowest node-metastability). The dashed red line delimits the 10 brain areas showing the highest node-metastability values for each group, respectively. When considering the top 10 nodes with the highest metastability in the HC group (0.102 ± 0.0011, mean ± s.d.), we found a consistent decline in both MCI (0.087 ± 0.0029) and AD(0.070 ± 0.0036), with values reduced across all corresponding Glasser parcels (*P*_FDR-corr_ *<* 0.0001). Importantly, some of these nodes (Area 7m, Area 7A and Area i6-8, see Table 2) are most likely to be rich-club hubs and positioned at the top of the cortical hierarchy. Overall, this pattern supports the hypothesis that hub regions may be particularly vulnerable to network degradation in AD progression.

**Table 2.** Top 10 Brain Regions Based on Meta-Analytic Estimates for Healthy Controls. Abbreviations: HC, Healthy Controls; MCI, Mild Cognitive Impairment; AD, Alzheimer’s Disease, Node-meta: node-metastability, L: left, R:right.

| Glasser Label | Node-meta HC | Node-meta MCI | Node-meta AD | Full name |
| --- | --- | --- | --- | --- |
| L_7AL_ROI | 0.103 | 0.087 | 0.076 | Lateral Area 7A |
| L_7m_ROI | 0.102 | 0.087 | 0.071 | Area 7m |
| L_i6-8_ROI | 0.102 | 0.081 | 0.066 | Inferior 6-8 Transitional Area |
| L_IP0_ROI | 0.104 | 0.089 | 0.071 | Intraparietal sulcus |
| L_LBelt_ROI | 0.101 | 0.086 | 0.066 | Lateral Belt Complex |
| L_TF_ROI | 0.101 | 0.087 | 0.071 | Area TF |
| R_33pr_ROI | 0.102 | 0.092 | 0.064 | Area 33 prime |
| R_FOP4_ROI | 0.102 | 0.092 | 0.074 | Frontal Opercular Area 4 |
| R_Ig_ROI | 0.104 | 0.086 | 0.069 | Insular Granular Complex |
| R_VMV2_ROI | 0.103 | 0.087 | 0.072 | VentroMedial Visual Area 2 |

**Table 3.** Machine Learning: Evaluation of Performance. Abbreviations: Nf, number of features; k, number of neighbors in the best kNN classifier; NA, not applicable (see Methods).

| Model | Accuracy (test,%) | AUC (test,%) | Accuracy (train,%) | AUC (train,%) | Best k | Nf |
| --- | --- | --- | --- | --- | --- | --- |
| HC vs AD | 85.71 | 95.83 | 90.0 | 97.80 | 4 | 10 |
| HC vs MCI | 71.43 | 65.0 | 84.21 | 98.21 | 2 | 10 |
| Overall | 78.57 | 80.41 | 87.10 | 98.0 | NA | 21.3 |

To further quantify the deviation from the cortical hierarchy, we calculated the hierarchy disruption factor (HDF). Our HDF measure showed a clear difference when comparing the cohort-averaged values: *HDF*(*HC, MCI*) = 0.13 and *HDF*(*HC, AD*) = 0.46. However, this difference did not reach statistical significance, probably due to the high variability exhibited by the individual measures.

### 3.4 MCI and AD stages are characterized by a reduction in node-metastability across RSNs

Next we examined group differences in node-metastability across RSNs, i.e., default (DMN), limbic (LIM), control (CNT), dorsal attention (DAT), visual (VIS), salience (SAL) and somatomotor (SM). Differences between groups for each RSN are depicted in **Figure 3A**. Compared to HC, the MCI group presented a significant decrease in node-metastability across all RSNs (*P*_FDR-corr_ *<* 0.001) except in the VIS network, which remained almost unchanged (*P*_FDR-corr_ *>* 0.05). Furthermore, the AD group showed a significant decrease in node-metastability across all RSNs compared to HC (*P*_FDR-corr_ *<* 0.001). Similarly, when compared to MCI, the AD group displayed a node-metastability decline in all networks, i.e., the CNT, SM, SAL, DMN, and DAT (*P*_FDR-corr_ *<* 0.001,) and LIM (*P*_FDR-corr_ *<* 0.05). **Figure 3B** shows a radar plot with a fingerprint of each group’s average node-metastability values for each RSN.

**Figure 3.**
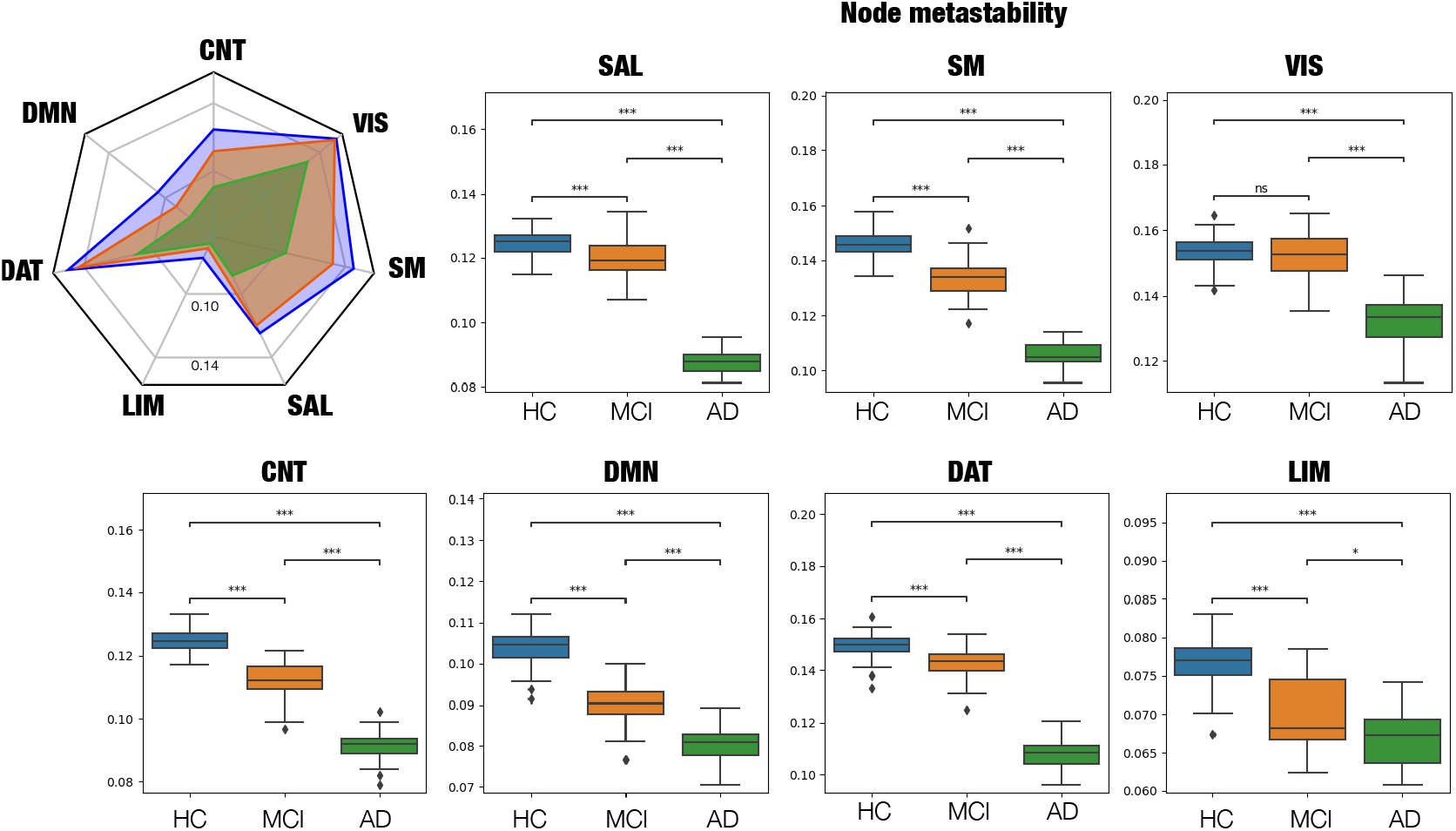
Node-metastability within resting state networks (RSNs). The radar plot (upper left) represents the average node-metastability values per RSN for each stage. Compared to HC subjects, node-metastability significantly decreased in the MCI and AD stages for all RSNs. P-values are based on a permutation test, where ns denotes non-significant and *** denotes *p <*= 1.00*e−* 03.

### 3.5 Node-metastability is linked to proteinopathy

As a confirmatory analysis, we first quantified Amyloid-*β* (A*β*) and tau burdens for each diagnostic group **Figure 4**. As expected, both protein burdens exhibited a systematic increase with disease progression, reaching statistical significance for the HC-AD comparison in both proteins and for the HC-MCI comparison in tau levels.

**Figure 4.**
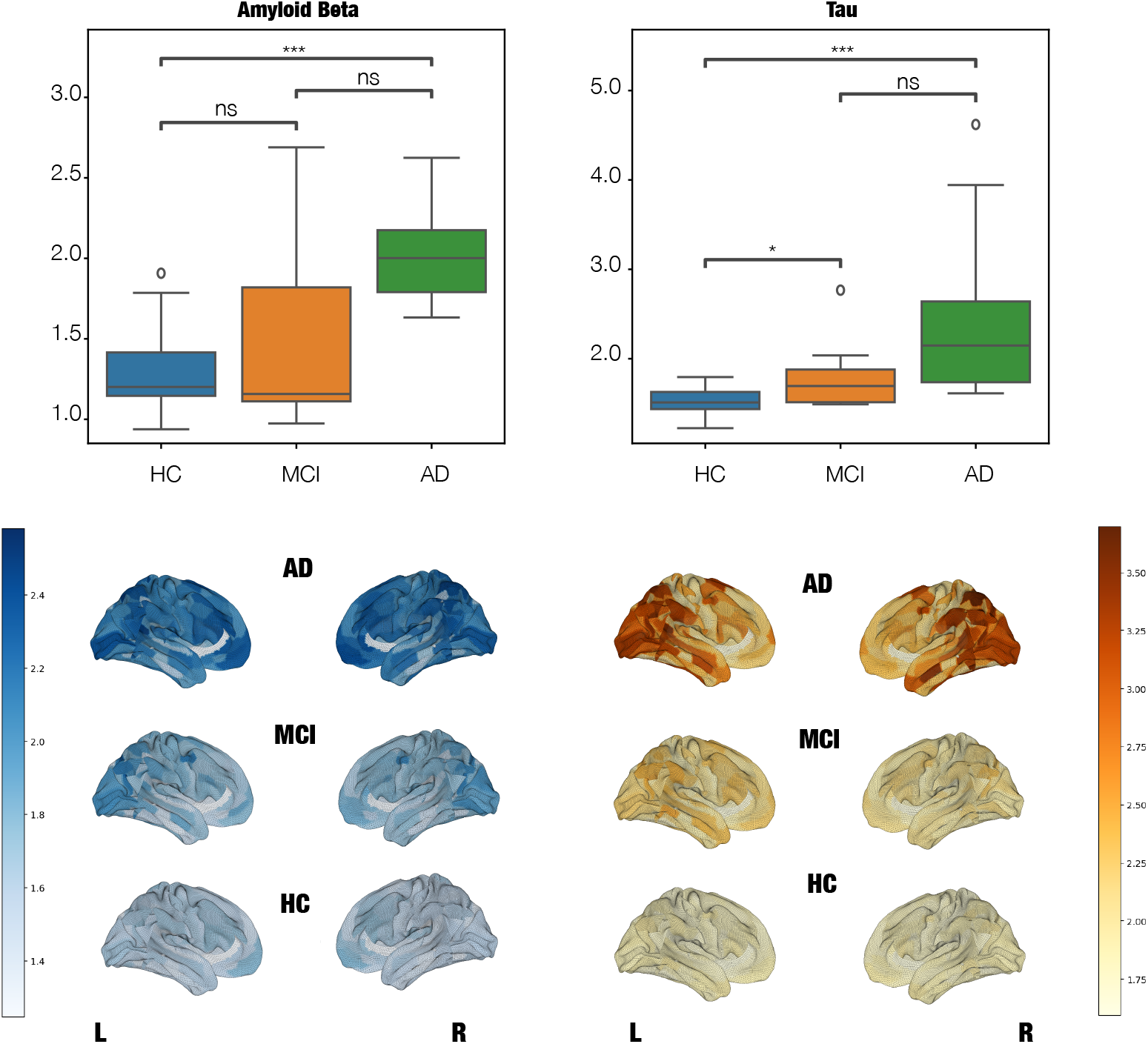
Average per-subject levels of Amyloid-*β* and Tau for the different AD stages. Top panel shows statistical differenes across AD groups. There is a significant increase of both burdens at advanced AD stages. P-values are based on a permutation test, where ns denotes non-significant, * denotes *p <*= 5.00*e−* 02, ** denotes *p <*= 1.00*e−* 02 and *** denotes *p <*= 1.00*e−* 03. Bottom panel shows brain renders showing the anatomical distribution of Amyloid-*β* and Tau for the different AD stages.

We then used mixed effects models to investigate the impact of both burdens, A*β* and tau, on node-metastability. The metastability for each node was defined as the outcome variable, with each region’s A*β* and tau SUVR values included as fixed effects. Subject ID was modeled as a random effect. We studied two different but complementary models: one assessing the influence of both burdens on node-metastability at the whole-brain level and the second one including the node-metastability of each RSN.

At the whole brain level, we observed a clear accumulation of A*β* and tau in the brain as the disease progresses, with node-metastastability significantly decreasing with both burdens **(Figure 5A-B**. The analysis shows a significant dependence of node-metastability on A*β* (Estimate = 1.662e-03, Std. Error = 6.766e-04, *P* = 0.014). In this analysis, tau and the interaction between the proteins did not reach statistical significance.

**Figure 5.**
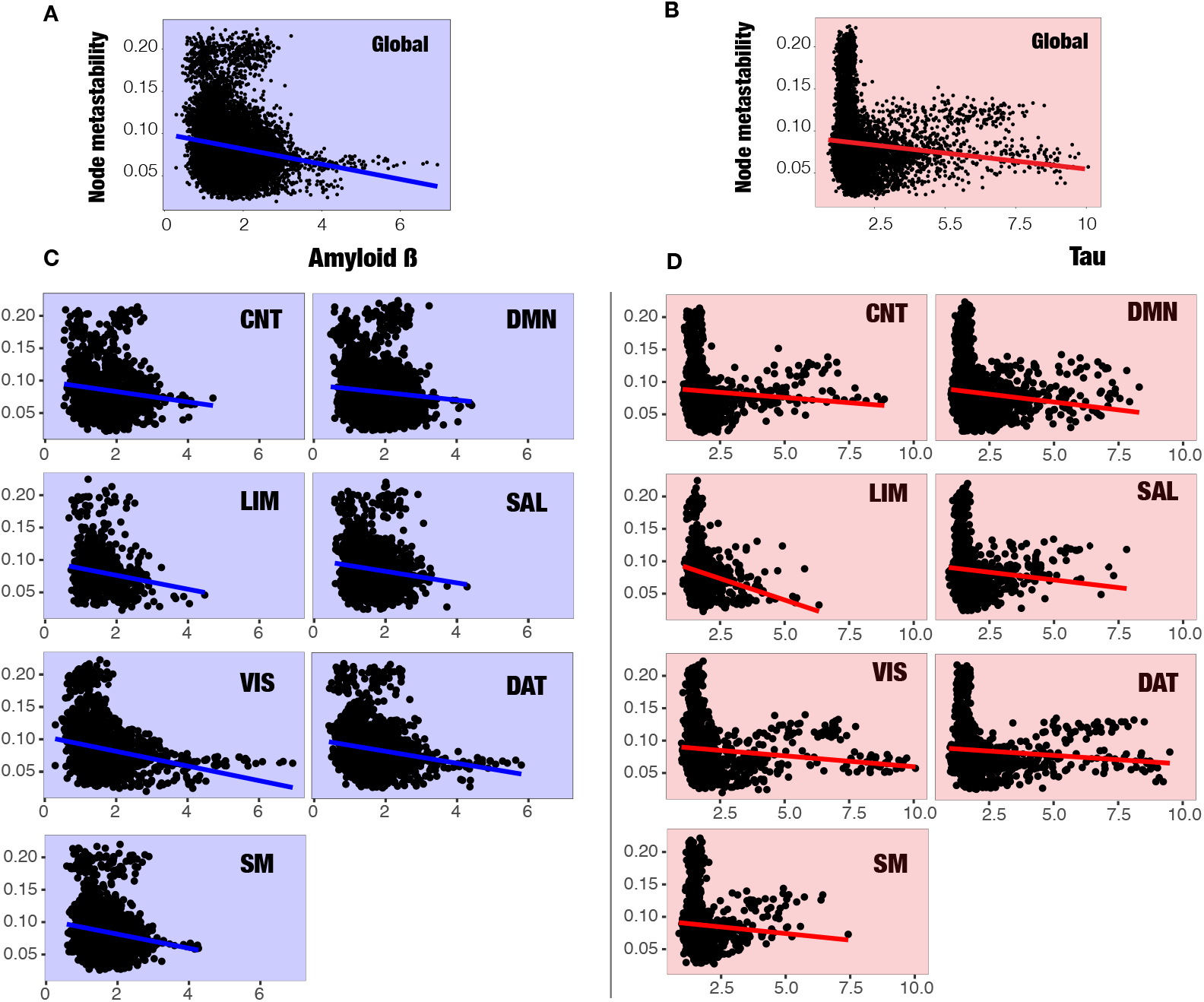
Multilevel models of the effects of Amyloid-*β*. **(A***β*) and Tau on whole-brain metastability (top panels) and for each RSN (bottom panels). Points represent the observed data, and lines represent the multilevel model-implied intercepts (mean of model coefficients for the random effect of subjects) and slopes (model coefficients for A*β* and tau). (A) We can observe the effect of A*β* (in blue) on node-metastability in the whole brain. From the model, we know that the effect of A*β* on node-metastability is significant (p = 0.014 *). (B) In red, we have the effect of Tau on node-metastability in the whole brain, which did not yield statistically significant results. (C) Here we plot the multilevel model-implied intercepts and slopes of the A*β* on metastability (blue), which did not yield significant results. (D) The effect of tau (red) is significant for both the DAT network (p = 0.00819 **) and the SM network (p = 0.04699 *) but not on the other networks. Finally, although not plotted, our analysis showed that the A*β* -tau interaction terms have a direct impact mainly on DAT network (p = 0.01348 *), showing the potential toxic feedback loop between both burdens on the disease evolution^51^.

By contrast, in the complementary RSN analysis, the role of tau was significant for both the DAT network(Estimate = 6.242e-03, Std. Error = 2.339e-03, *P* = 0.00819) and the SM network (Estimate = 7.007e-03, Std. Error = 3.519e-03, *P* = 0.04699). Although the effect of A*β* did not reach statistical significance**Figure 5C**, we found that the A*β* -tau interaction term has a direct impact mainly on the DAT Network (Estimate = -1.978e-03, Std. Error = 7.970e-04, *P* = 0.01348), showing the synergistic effect of both burdens on the disease evolution^51^ **Figure 5D**.

### 3.6 Machine learning-based classification of dementia stage

Finally, we investigated whether brain dynamics and protein burden would allow accurate classification of the dementia stage. To this end, we used independent k-nearest neighbors (kNN) classifiers to dissociate the diagnostic groups in this study (HC vs MCI and HC vs AD) based on brain dynamics (ignition, node-metastability) and proteinopathy (A*β* and tau SUVR) from each Glasser parcel (1440 features in total per participant). Minimal-optimal features were selected by an mRMR algorithm. To allow for comparisons across analyses, the number of selected features by the mRMR algorithm was fixed to 10 following prior literature ^52^. The number of neighbors was optimized in the range between 2-5 neighbors. Leave-one-out cross-validation was used to evaluate the performance of the model **Figure 6A**.

**Figure 6.**
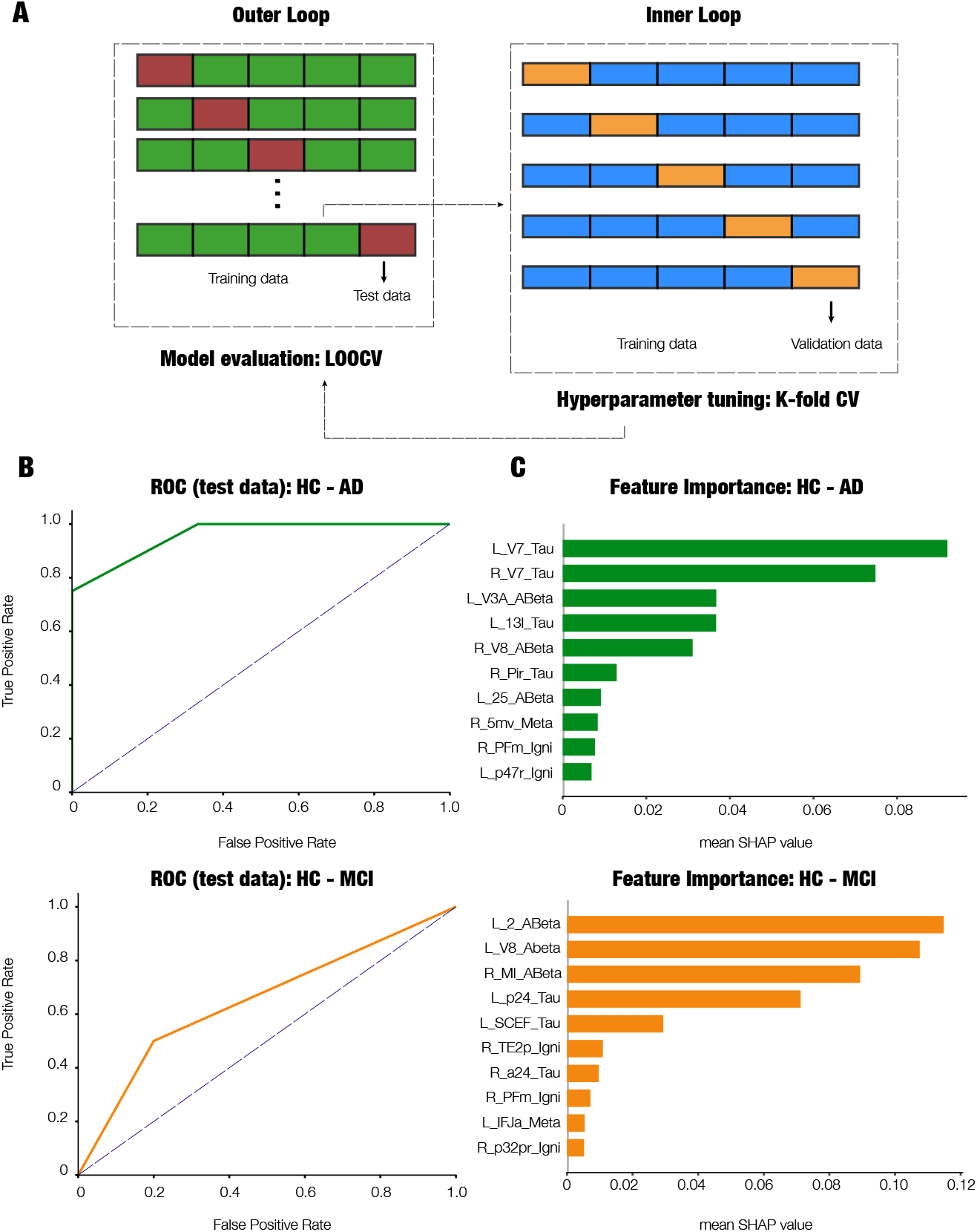
Machine learning classification of dementia stages based on brain dynamics and protein burden. (A) Overview of the nested cross-validation approach. A leave-one-out cross-validation was applied in the outer loop for model evaluation while a k-fold cross-validation for hyperparameter turning was performed in the inner loop. (B) ROC curves for test data. (C) 10 selected features selected by the mRMR algorithm rank by their predictive power on the model performance based on SHAP values. Abbreviations: L, left; R, right; ABeta, amyloid-*β* ; Meta: node-metastability, Igni, ignition. The corresponding Glasser parcel for each feature appears abbreviated as in the original publication ^37^.

The classification yielded predictions of diagnostic categories of 78.6 and 87.1% (mean accuracy across all binary problems) using training and test sets, respectively (Table 3). The ROC curves endorsed the between-categories discriminative ability of the classifiers, with a mean AUC as high as 80.4 and 98.1% for training and test data, respectively (Table 3, **Figure 6B**).

We used SHAP values to indicate the importance of each feature for classification. Tau burden was the most relevant feature in the classification of AD versus HC. In contrast, the most relevant feature for the classification of MCI from HC was A*β* burden (**Figure 6C**). This is in agreement with previous findings ^28^ using this dataset, where tau levels had a greater influence on more advanced stages (AD) of the disease, whereas A*β* were more important at earlier stages (MCI).

## 4 Discussion

In this study, we implemented a recently developed framework to characterize disruptions in how local activity influences global computation across the AD continuum using cross-sectional data from ADNI3. Specifically, we calculated intrinsic ignition and node-metastability measures, which reflect the capability of a given brain area to propagate neuronal activity to other regions and, importantly, how this propagation varies over time. As a result, node-metastability can provide insights into the exact nature of hierarchical information processing. Here, at the whole-brain level, we found a progressive and significant decline in node-metasbility at MCI and AD stages when compared to HC, suggesting a disruption of the cortical hierarchy information processing. Similar reductions were found across nearly all resting-state (RSNs) networks at MCI and AD stages. To link these alterations in large-scale brain dynamics to molecular hallmarks of AD, we also assessed the impact of amyloid beta (A*β*) and tau burden on brain dynamics. We found that both significantly impact the whole-brain functional network and, in general, all RSNs. Finally, we conducted a machine learning classification of dementia stages using brain dynamics and protein burden features and corroborated previous findings from a previous computational modelling study with this subset of participants ^28^, indicating a dominant role of A*β* and tau burden at MCI and AD stages, respectively.

At the whole-brain network, the dynamical complexity decreased progressively according to diagnostic group, with AD patients showing the lowest values, indicating a gradual deterioration in brain network dynamics **(Figure 2A)**. Only a few works have analyzed the impact of AD on whole-brain dynamics and information processing across large-scale brain networks. For instance, Sanz-Arigita et al.^53^ did a graph analysis of the resting-state fMRI functional connectivity (rs-FC) and found that the empirical data pointed to increased synchronization of frontal cortices, together with a clear decrease at the parietal and occipital areas, which results in a net reduction of functional long-distance links between frontal and caudal brain regions. Wu et al.^54^ found similar results using group information-guided ICA, showing that rs-FC alterations mostly appear in the temporal, cingulate, and angular areas with a pattern consistent with regions implicated in brain atrophy ^55^ and A*β* deposition ^56^ in AD. Taken together, these studies support the notion that AD-related neuropathology disrupts the dynamical integrity, in terms of node-metastability, of large-scale brain networks.

The human connectome is organized into a densely interconnected core of high-degree hub nodes known as the rich-club ^57^. Here, we found that AD progression is associated with a systematic decrease in the cortical hierarchy of information processing that could explain the diminished global integration of driving events. This suggests that node-metastability may be a signature of disruption in the rich-club organization. Our result aligns with previous literature showing that AD is associated with decreased metastability and disrupted rich-club organization ^20^. In support of this notion, some of the nodes positioned at the top 10 of the cortical hierarchy, as measured with node-metastability, were most likely to be rich-club hubs (Table 2). Specifically, Area 7m in the posterior parietal cortex emerged as a prominent candidate, aligning with its established role as a core node of the DMN. It exhibits high-degree centrality, extensive long-range connectivity, and characteristically low myelination — features consistent with its placement at the apex of the cortical processing hierarchy ^58^. Functionally, 7m is critically involved in internally directed cognition and episodic memory, and it has been consistently classified as a rich-club hub in both structural and functional connectomic analyses ^57^. Lateral Area 7A also shows strong attributes of a high-level integrative node. As part of the DAT, it interfaces with frontal eye fields and intraparietal areas, supporting visuospatial integration and attentional control ^59^. Of note, as later discussed, we also found tau accumulation and A*β* ×tau at the DAT were negatively associated with node-metastability.

Recent resting-state fMRI has enabled exploring the brain’s intrinsic organization of large-scale distributed networks, revealing that AD modulates brain dynamics in RSNs, including a strong reduction in rs-FC in the DMN, SAL and subcortical networks during the initial stages^11^–19. Our results revealed node-metastability disease-dependent changes across large-scale RSNs **(Figure 2)**. In particular, we found that the VIS network exhibited the highest node-metastability values during all stages, remaining relatively unchanged in MCI to HC but showing a severe decrease in later AD stages. In general, we observe a significant reduction in the node-metastability of all RSNs along the disease progression, except for the VIS network. This finding is interesting given that the VIS network is the one that exhibits the highest structure-function decoupling index and therefore might be more resilient to the structural damage caused by the disease ^60^. However, future studies should confirm this interpretation.

In line with prior evidence linking proteinopathy progression to network-level dysfunction in AD ^61–63^, our findings revealed a burden-dependent reduction in node-metastability associated with A*β* and tau accumulation. Mixed-effects models at the whole-brain level identified a significant inverse relationship between global node-metastability and A*β* burden, whereas tau and A*β* ×tau interaction terms did not reach significance, suggesting that A*β* alone might exert a widespread destabilizing effect on functional network dynamics in early disease stages. However, when restricting the analysis to RSNs, a more nuanced pattern emerged: tau pathology showed significant associations with reduced node-metastability in both the DAT and SM networks, highlighting regional vulnerabilities to tau-mediated network disruption. Moreover, we observed a significant A*β* ×tau interaction effect within the DAT network, supporting a synergistic contribution of these hallmark pathologies to RSN-specific impairments in node-metastability. This result aligns with preclinical models suggesting that A*β* -induced neuronal hyperactivity might potentiate tau-related network collapse in a regionally selective manner, particularly in higher-order associative networks implicated in early AD pathogenesis ^51^. Together, these findings extend previous work by identifying differential and interacting roles of A*β* and tau in destabilizing large-scale brain dynamics.

Here, we also demonstrated that measures of brain dynamics—specifically ignition and node metastability—alongside protein burden (A*β* and tau), enable accurate classification of individuals with MCI and AD. Consistent with findings from a previous computational modeling study^28^, A*β* burden emerged as the most accurate feature for classifying individuals with MCI. In contrast, tau levels provided the highest classification accuracy for patients with AD. Notably, none of the brain dynamics features included in the model ranked among the top in feature importance. Future studies including participants with subjective cognitive decline (SCD) might help clarify whether brain dynamics serve as more sensitive biomarkers for early stages, as suggested by a recent MEG study ^52^. If confirmed, dynamic brain measures derived from fMRI could contribute to the triaging of patients in the early stages of AD, particularly when clinical diagnosis is complicated by high cognitive reserve ^64^.

We acknowledge several limitations in the present study. First, the findings are based on a relatively small cohort, and future studies should aim to replicate these results in larger samples with balanced representation across diagnostic groups. Second, the cross-sectional design precludes causal inference; the inclusion of longitudinal data in future research could provide stronger evidence regarding the temporal dynamics of the observed effects. Third, we employed the Glasser parcellation scheme to ensure comparability with previous studies using this dataset; however, the robustness of our findings should be assessed using alternative parcellation approaches in future work. Finally, additional metrics of brain dynamics — such as turbulent dynamics, which have demonstrated high classification accuracy in other neuropsychiatric populations^65^ — were not examined in this study and warrant investigation in future analyses.

In summary, our findings have significant implications for understanding the alterations in brain dynamics across the progression of AD. This study demonstrates that disease stage influences the dynamic complexity of both the whole-brain functional network and large-scale RSNs, as well as the cortical hierarchy involved in information processing. Furthermore, the associations between brain dynamics and toxic protein burden suggest that, when combined with computational modeling, these measures could contribute to the prediction of treatment efficacy. In particular, this may apply to pharmacological and other therapeutic interventions aimed at reducing pathological protein accumulation, potentially mitigating the disruptions in brain dynamics at various stages of AD progression as observed in the present study.

## Data and code availability statement

Upon acceptance, all code for implementing computational models and reproducing our results will be available at https://github.com/dagush/WholeBrain

## Ethics Statement

Ethical approval was obtained by ADNI sites and written informed consent was collected from all participants. No further consent was necessary.

## Acknowledgements

We would like to thank our former master’s student, Lars Rass, for his invaluable assistance in developing the machine learning pipeline. Data collection and sharing for this project was funded by the Alzheimer’s Disease Neuroimaging Initiative (ADNI) (National Institutes of Health Grant U01 AG024904) and DOD ADNI (Department of Defense award number W81XWH-12-2-0012). ADNI is funded by the National Institute on Aging, the National Institute of Biomedical Imaging and Bioengineering, and through generous contributions from the following: AbbVie, Alzheimer’s Association; Alzheimer’s Drug Discovery Foundation; Araclon Biotech; BioClinica, Inc.; Biogen; Bristol-Myers Squibb Company; CereSpir, Inc.; Cogstate; Eisai Inc.; Elan Pharmaceuticals, Inc.; Eli Lilly and Company; EuroImmun; F. Hoffmann-La Roche Ltd and its affiliated company Genentech, Inc.; Fujirebio; GE Healthcare; IXICO Ltd.; Janssen Alzheimer Immunotherapy Research & Development, LLC.; Johnson & Johnson Pharmaceutical Research & Development LLC.; Lumosity; Lundbeck; Merck & Co., Inc.; Meso Scale Diagnostics, LLC.; NeuroRx Research; Neurotrack Technologies; Novartis Pharmaceuticals Corporation; Pfizer Inc.; Piramal Imaging; Servier; Takeda Pharmaceutical Company; and Transition Therapeutics. The Canadian Institutes of Health Research is providing funds to support ADNI clinical sites in Canada. Private sector contributions are facilitated by the Foundation for the National Institutes of Health (www.fnih.org). The grantee organization is the Northern California Institute for Research and Education, and the study is coordinated by the Alzheimer’s Therapeutic Research Institute at the University of Southern California. ADNI data are disseminated by the Laboratory for Neuro Imaging at the University of Southern California.

## Appendix: Collaborators

Data used in the preparation of this article were obtained from the Alzheimer’s Disease Neuro Imaging Initiative (ADNI) database (`adni.loni.usc.edu`). As such, the investigators within the ADNI contributed to the design and implementation of ADNI and/or provided data but did not participate in the analysis or writing of this report. A complete listing of ADNI investigators can be found at: http://adni.loni.usc.edu/wp-content/uploads/how_to_apply/ADNI_Acknowledgement_List.pdf

---

**Fig S1.**
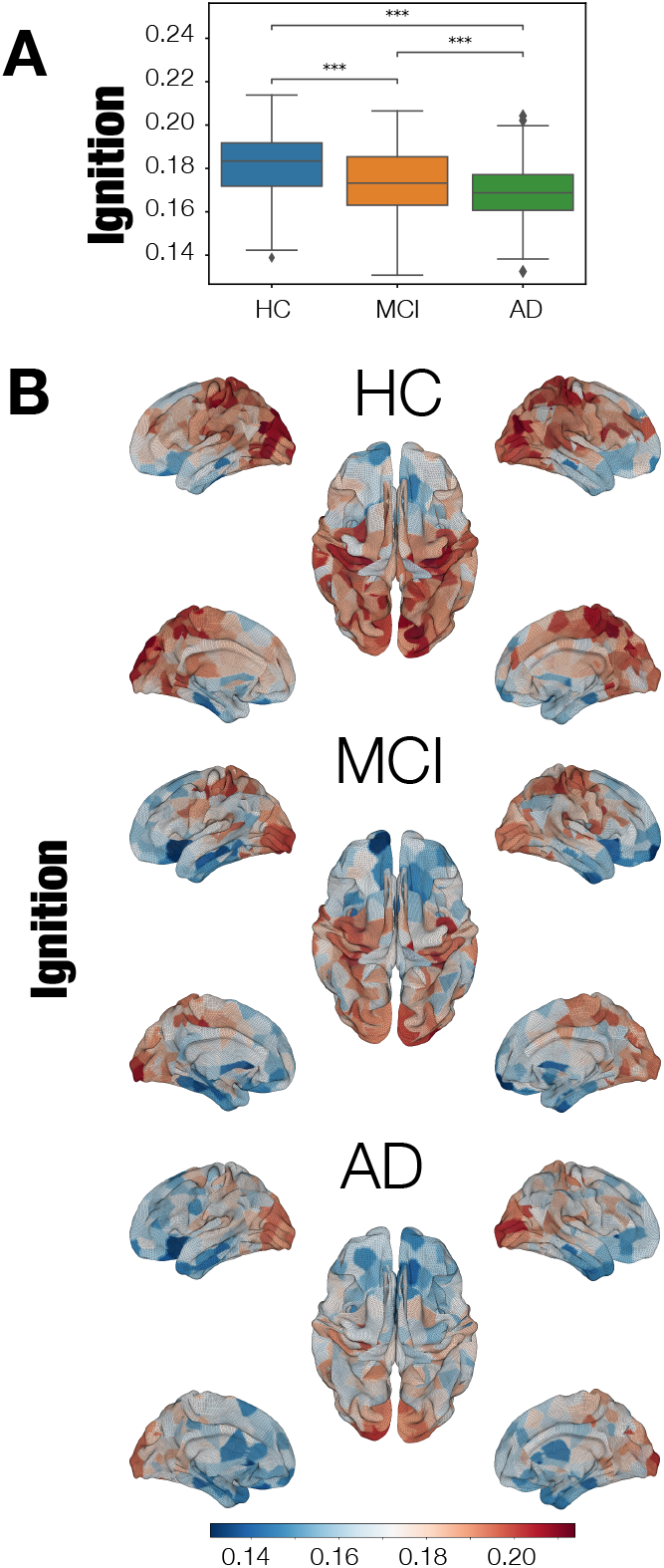
Intrinsic ignition is reduced across the AD continuum. **(A)** Ignition. HCs showed higher ignition values at the whole-brain level compared to MCI and AD patients. P-values are based on a permutation test, where *** represents *p <*= 1.00*e−* 04. **(B)** Brain renders represent the ignition values of the 379 areas for each AD stage.

